# Aramchol attenuates fibrosis in mouse models of biliary fibrosis and blocks the TGFβ-induced fibroinflammatory mediators in cholangiocytes

**DOI:** 10.1101/2024.11.06.621880

**Authors:** Sayed Obaidullah Aseem, Grayson Way, Jing Wang, Derrick Zhao, Yunling Tai, Emily Gurley, Jing Zeng, Xuan Wang, Phillip B Hylemon, Robert C. Huebert, Arun J. Sanyal, Huiping Zhou

**Author notes:** Contact: Sayed Obaidullah Aseem, M.D.,Ph.D., Stravitz-Sanyal Institute for Liver Disease & Metabolic Health, 1220 East Broad Street, Room 5036, Richmond, VA 23298.

## Abstract

**Background:** Fibroinflammatory cholangiopathies, such as primary sclerosing cholangitis (PSC) and primary biliary cholangitis (PBC), are characterized by inflammation and biliary fibrosis, driving disease-related complications. In biliary fibrosis, cholangiocytes activated by transforming growth factor-β (TGFβ) release signals that recruit immune cells to drive inflammation and activate hepatic myofibroblasts to deposit the extracellular matrix (ECM). TGFβ regulates stearoyl-CoA desaturase (SCD), an enzyme that catalyzes the synthesis of monounsaturated fatty acids, in stimulating fibroinflammatory lipid signaling. However, the role of SCD or its inhibitor, Aramchol, has not been investigated in biliary fibrosis or TGFβ-mediated cholangiocyte activation.

**Method:** 10–16-week-old multi-drug resistance 2 knockout (Mdr2^-/-^) and 3,5-diethoxycarboncyl-1,4-dihydrocollidine (DDC) diet-fed mice were orally gavaged daily with Aramchol at 12.5 mg/kg/day for 4 and 3 weeks, respectively. Liver and serum were harvested for the assessment of fibrosis and inflammation. Transformed human cholangiocyte cells (H69) and mouse large biliary epithelial cells (MLEs) were used to test the effects of the SCD inhibitor, Aramchol, at varying doses on TGFβ-mediated expression of fibroinflammatory signals and were confirmed in PSC-derived cholangiocytes (PSC-Cs) using ELISA, qPCR, and Western blot analyses.

**Results:** Aramchol treatment of Mdr2^-/-^ mice with established biliary fibrosis (treatment) and DDC diet-induced (prevention) models of cholestatic injury and fibrosis demonstrated significant reductions in both measures of ECM synthesis (mRNA expression of ECM components in the liver), collagen content of the liver (picrosirius red staining and hydroxyproline content) and myofibroblast activation (αSMA staining). *Il6* and *Tnfa* were also reduced with Aramchol in the liver. RNA-seq analysis of H69 cells showed that Aramchol co-treatment led to significant inhibition of TGFβ-induced hepatic fibrosis pathways while upregulating peroxisome proliferator-activated receptor (PPAR) signaling. *SCD* expression was significantly increased in TGFβ-treated H69 cells (2-fold, p<0.05). Aramchol in a dose-dependent manner significantly attenuated the increased expression of the fibrotic marker, plasminogen activator inhibitor-1 (PAI-1/SERPINE1), and hepatic stellate cell-activating genes (*VEGFA* and *PDGFB*) in TGFβ-activated H69 and MLEs. Aramchol also markedly reduced the expression of the inflammatory cytokine, interleukin 6 (IL6). SCD siRNA knockdown produced similar results in H69 cells. Furthermore, in PSC-Cs, the expressions of SCD, VEGFA and IL6 were significantly reduced with Aramchol. The expression of the anti-fibroinflammatory factors PPARα and -γ were modestly increased in cholangiocyte cell lines with increased expression of PPAR-responsive genes and increased nuclear binding of DNA PPAR response elements with Aramchol co-treatment compared to TGFβ only.

**Conclusion:** Aramchol, an SCD inhibitor, both attenuates and prevents biliary fibrosis in mouse models of cholestatic injury and fibrosis. This effect is partially due to Aramchol inhibiting TGFβ-induced fibroinflammatory mediators in cholangiocytes by upregulating PPARα and -γ expression and activity. These findings, along with Aramchol’s excellent safety profile in clinical trials, provide the rationale for assessing Aramchol in further clinical studies in patients with biliary fibrosis, particularly PSC, where a treatment is desperately needed.

## INTRODUCTION

Biliary fibrosis is the predominant pathological process in fibroinflammatory cholangiopathies, notably primary sclerosing cholangitis (PSC) and primary biliary cholangitis (PBC). These diseases collectively account for at least 16% of liver transplantation performed in the US with an annual cost of $400 million [1]. Yet, there are no approved treatments for biliary fibrosis and late stage cholangiopathies typically require liver transplantation.

A key pathway of fibrosis is transforming growth factor beta (TGFβ) signaling, a potent activator of hepatic stellate cells and portal fibroblasts into a myofibroblast-like phenotype that secretes extracellular matrix (ECM) [2]. Immune cells such as macrophages and cholangiocytes are the major sources of secreted TGFβ [3, 4], which by autocrine or paracrine mechanisms stimulate cholangiocytes themselves. Cholangiocytes activated by TGFβ recruit transcriptomic regulators, including epigenetic enzymes, which result in the expression of fibroinflammatory signals [5-7]. Activated cholangiocytes secrete TGFβ, inflammatory mediators such as interleukin (IL) 6, and other fibrogenic factors such as plasminogen activator inhibitor 1 (PAI-1/SERPINE1), further propagating the fibroinflammatory signaling [8-10].

TGFβ also regulates cellular metabolic and lipid profile [11, 12]. However, how these cellular metabolic changes enable TGFβ-mediated signaling in biliary fibrosis is not completely elucidated. In epithelial cells, TGFβ upregulates the expression of stearoyl-CoA desaturase-1 (SCD1) [11-13], an enzyme that catalyzes the rate-limiting step in the formation of mono-unsaturated fatty acids (MUFAs), notably palmitoleate (C16:1n-7) and oleate (C18:1n-9) from palmitic acid (C16:0) and stearic acid (C18:0). SCD1 substrates and products have demonstrated both pro- and anti-inflammatory effects [14-17]. Additionally, SCD1 suppresses peroxisome proliferator-activated receptor (PPAR)-γ in various cells [11, 13]. These observations suggest cell-specific signaling and require further investigation of SCD1 in cholangiocytes.

Arachidyl-amido cholanoic acid (Aramchol) was previously shown to both downregulate SCD1 and upregulate PPARγ expression. Aramchol has demonstrated anti-fibrotic effects in hepatic stellate cells (HSCs) and hepatocytes, thereby regulating markers of fibrosis in mouse models of metabolic dysfunction-associated steatotic liver disease (MASLD) [18-20]. In clinical trials of MASLD, Aramchol demonstrated beneficial effects on steatohepatitis and fibrosis with an excellent safety profile [21, 22]. However, the role of Aramchol in cholangiocyte signaling and biliary fibrosis was not previously assessed. In this study, we demonstrate that Aramchol significantly inhibited fibrosis and inflammatory markers in two mouse models of biliary fibrosis. Furthermore, we showed that Aramchol inhibited the TGFβ-stimulated fibroinflammatory signals in cholangiocytes while upregulating PPARα/γ activity. Thus, this study provides the rationale for assessing Aramchol in clinical trials of human diseases driven by biliary fibrosis, particularly PSC.

## MATERIALS AND METHODS

### Mouse studies

Multi-drug resistance 2 knockout (Mdr2^-/-^) mice on C57BL/6J background were kindly provided by Dr. Daniel Goldenberg (Hadassah-Hebrew University Medical Center, Jerusalem, Isreal). For *in vivo* studies, the salt form of Aramchol, Aramchol meglumine, was used. Male and female 10–16-week-old Mdr2^-/-^ mice were administered Aramchol meglumine dissolved in water at 12.5 mg/kg or vehicle only by oral gavage daily for four weeks. At the end of the experiment, mice were euthanized, and liver and blood harvested.

Similarly, wild-type female and male mice were placed on regular chow or 3,5-diethoxycarboncyl-1,4-dihydrocollidine (DDC) diet for 3 weeks with 1-day breaks of regular chow every week. All mice were orally gavaged with Aramchol meglumine or vehicle as described above.

All mice were housed under 12-12-hour light and dark cycles with water and standard chow *ad libitum* when not on DDC diet. All mouse experiments were conducted in strict accordance with protocols approved by the Virginia Commonwealth University Institutional Animal Care and Use Committee.

### Hydroxyproline measurement

Hydroxyproline content of mouse livers were quantified using mass spectrometry by VCU Lipidomics and Metabolomics core. Briefly, area under the curve of hydroxyproline standards were used to generate a standard curve, which was used to measure the concentration of samples. Sample measurements were normalized to their weights (30-50 mg).

### RNA-sequencing

RNA sequencing and bioinformatics analysis was conducted with the VCU Genomics and Bioinformatics cores. RNA quality control was performed using Agilent Bioanalyzers. Pathway analyses were performed by Ingenuity Pathway Analysis (IPA, Qiagen).

### Cell culture

Transformed cholangiocyte cell line, H69 cells, were cultured in Dulbecco’s Modified Eagle Medium (DMEM)/F12 supplemented with 10% fetal bovine serum, 1% penicillin/streptomycin, adenine, insulin, epinephrine, T3-T, hydrocortisone, and epidermal growth factor. PSC patient-derived cholangiocytes (PSC-Cs) were kindly provided by Dr. Nicholas LaRusso (Mayo Clinic, Rochester, MN, USA) and cultured in the H69 media [23, 24]. Mouse large biliary epithelial cells (MLEs) were cultured in complete DMEM media [25, 26]. Cells were serum starved before treatment with 10 ng/mL recombinant TGFβ (R&D Systems #240-B) for 16 hours. For siRNA targeting, cells were transfected with SCD1 or control siRNAs (Dharmacon, ON-TARGETplus SMARTPool) using the Oligofectamine reagent (Invitrogen #12–252-011). Aramchol acid (Galmed Pharmaceutical) was dissolved in dimethyl sulfoxide (DMSO) first and cells treated at the indicated doses. For some experiments, nuclear extracts were isolated using a nuclear isolation kit (Cayman Chemical, Cat# 10009277). Nuclear extracts were used to show PPARγ and PPARα binding to PPAR responsive elements (PPRE) using colorimetric kits (Abcam ab133107; Cayman Chemical Cat# 10006855).

### Quantitative RT-PCR

Total RNA was extracted from cells using the TRIzol RNA Isolation Reagent (ThermoFisher, 15596026). Reverse transcription was performed with 2μg RNA using oligo (dT) primer and SuperScript III. Real-time PCR was performed using Sybr Green Master Mix made to 20 μl final volume per reaction and the QuantStudio™ 3 Real-Time PCR System (Applied Biosystems™) using primer sequences shown in Supplemental Table 1.

### ELISA

Mouse and human IL6 ELISA kits with standards were purchased from eBioscience (Human Cat# 88-7066-88, Mouse Cat# 88-7064-88). IL6 was quantified in the cell cultured media following manufacturer protocols.

### Immunoblotting

Cells were lysed, and protein isolated in RIPA buffer. Protein concentration was determined using BCA protein assay. Cell cultured media were briefly span down before immunoblotting. 10-40μg of protein was loaded onto either 4–20% or 10% Tris-Glycine gels, electrophoresed, and transferred onto nitrocellulose membranes (Scientific Laboratory Supplies) for blotting. The membrane was blocked, incubated with primary antibodies, rinsed with TBST, and incubated with fluorophore conjugated secondary antibodies (Supplemental Table 1). After washing, the membrane was imaged using iBright 1500 imaging system (Invitrogen).

### Immunofluorescence and imaging

Cells were fixed with formalin for 15 minutes. De-paraffinized liver tissue or cell slides were blocked using Intercept Blocking Buffer (LI-COR #927-70001) followed by overnight incubation with the primary antibody followed by species specific fluorophore conjugated secondary antibodies. Slides were imaged with Axio Observer A1 inverted (Zeiss) and Keyence BZ-X910 microscopes. Mouse liver slides were imaged at 10X and stitched together for further analysis using ImageJ software for area quantification.

### Statistical analysis

Data are presented as mean with standard deviation. One-way analysis of variance (ANOVA) with Tukey’s Honestly Significant Difference (HSD) post-test was used to assess the statistical significance between ≥3 groups with P-value <0.05 in two-tailed analyses. Where data did not have a normal distribution, one-way ANOVA on ranks (Kruskal Wallis Test) with post-hoc Dunn’s test was used. Student’s t-test was used in 2 group analyses.

## RESULTS

### Aramchol meglumine downregulates fibrosis and inflammation in mouse models of biliary fibrosis

Aramchol reduced steatosis and fibrosis in a dietary mouse model of MASLD [20]. We assessed the effect of Aramchol on fibrosis and inflammation in 2 mouse models of biliary fibrosis. In the first, Mdr2^-/-^ mice, which have established fibrosis and inflammation at 10–16-weeks of age, were used as a reversal (treatment) model. The Aramchol meglumine formulation at 12.5 mg/kg/day or vehicle (water) only was administered daily for 4 weeks. Overall, the mice did not show any signs of toxicity and tolerated the drug well. There were no differences in weight gain (Supplementary Fig. 1A) or behavioral changes. Additionally, there were no significant changes observed in liver chemistries, including serum hepatic injury markers (alkaline phosphatase (ALP), alanine aminotransferase (ALT), and aspartate aminotransferase (AST)) or hepatic function tests (total bilirubin (TB) and albumin) (Supplementary Fig. 1B). Aramchol meglumine significantly reduced fibrosis as demonstrated by picrosirius red staining and quantification of area of positive staining (Fig. 1A and B). Consistently, the mRNA expression of extracellular matrix content, Collagen (*Col1a*) and α-Smooth muscle actin (α*Sma*) were significantly reduced (Fig. 1C). Similarly, pro-fibrotic markers PAI1/Serpine1 and Timp1 were significantly reduced with Aramchol meglumine treatment (Fig. 1C). Markers of inflammation, *Il6, Tnfa* and *Nfkb*, were also significantly reduced with Aramchol meglumine (Fig. 1D). Hydroxyproline content of the liver trended down with Aramchol meglumine treatment but did not reach statistical significance (p=0.09). Myofibroblast activation was significantly reduced with Aramchol meglumine treatment as demonstrated by immunofluorescence using αSMA marker (Fig. 1F).

**Fig 1.**
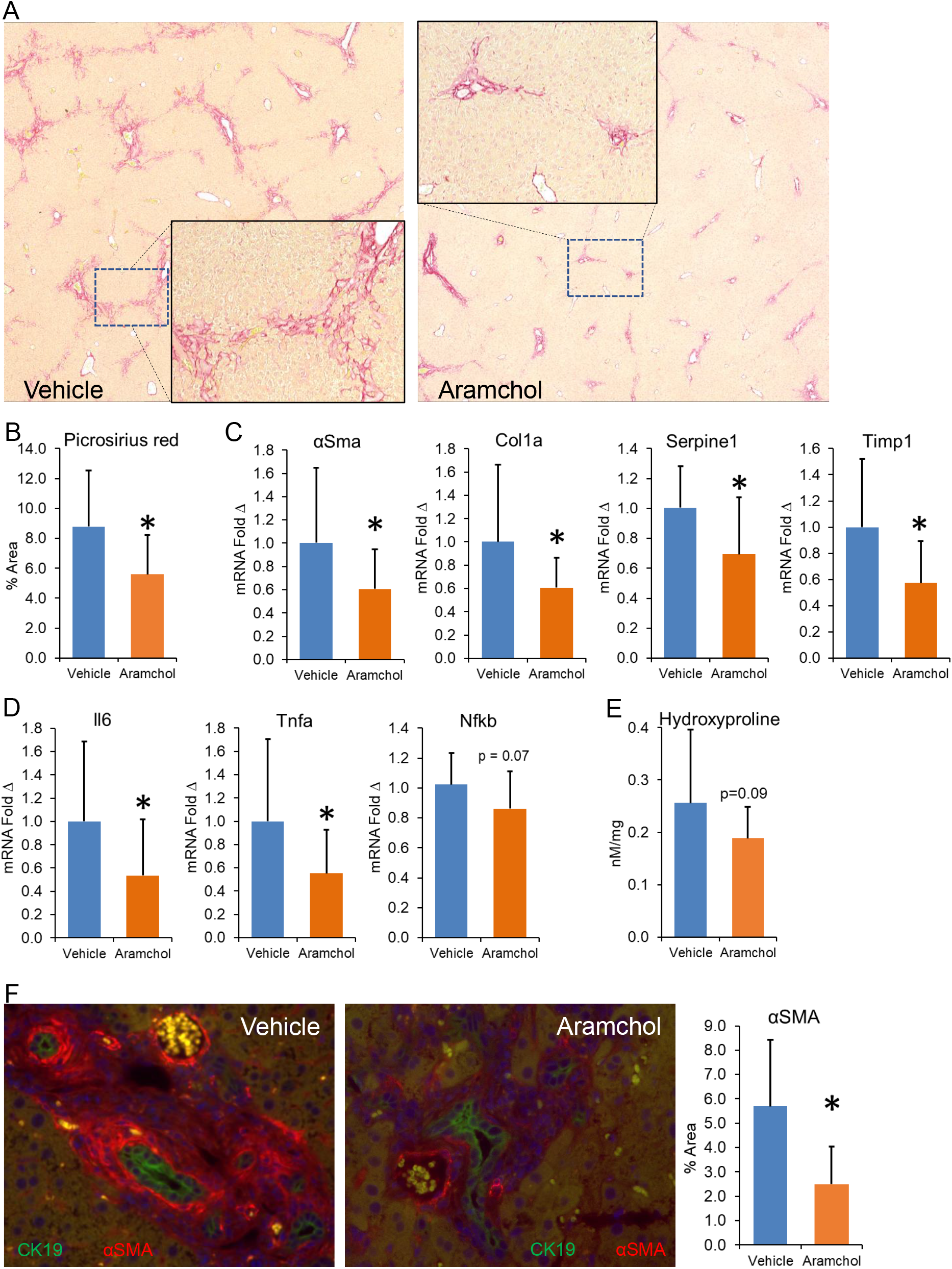
A) Representative images of picrosirius red staining of livers from Mdr2^-/-^ mice showed reduced staining with Aramchol meglumine treatment (shortened to Aramchol in the figures) compared to vehicle with area quantification using ImageJ shown in B. C&D) qPCR analysis of liver lysate demonstrated attenuation of ECM components, collagen (*Col1a*) and alpha smooth muscle actin (*αSMA*), pro-fibrotic markers *Serpine1* and *Timp1*, and inflammatory markers, *Il6* and *Tnfa*. E) Hydroxyproline content in the liver trended down with Aramchol meglumine treatment compared to vehicle but did not reach statistical significance (p=0.09). F) Immunofluorescence (IF) of αSMA (red) showed reduced immunostaining surrounding CK19 immunostained bile ducts (green) with Aramchol meglumine treatment. (N=9 mice per group, *= p<0.05).

In the second fibrosis prevention model, 10-16-week-old, age and sex matched wild type mice were placed on the DDC diet with 1-day break of regular chow weekly for 3 weeks. Mice received Aramchol meglumine or vehicle daily. DDC diet resulted in a significant decrease in weight but there was no significant difference between the Aramchol meglumine and vehicle groups (Supplementary Fig. 2A). Similarly, there were no significant differences in serum ALP, ALT, AST, TB, or albumin in Aramchol meglumine treated mice compared to vehicle only (Supplementary Fig. 2B). The increase in fibrosis was significantly attenuated with Aramchol meglumine administration as demonstrated by significantly reduced hydroxyproline content (Fig. 2A). Consistently, the mRNA expressions of *Col1a*, α*Sma* and *Serpine1* were significantly reduced (Fig. 2B). Similarly, picrosirius red staining and quantification of positively stained area was significantly reduced with Aramchol meglumine treatment (Fig. 2C and D). The DDC diet-induced markers of inflammation, *Il6, Tnfa* and *Nfkb*, were similarly significantly reduced with Aramchol meglumine treatment (Fig. 2E). Myofibroblast activation determined by αSma staining of liver sections by immunofluorescence was significantly reduced in Aramchol meglumine treated mice compared to vehicle only (Fig. 2F).

**Fig 2.**
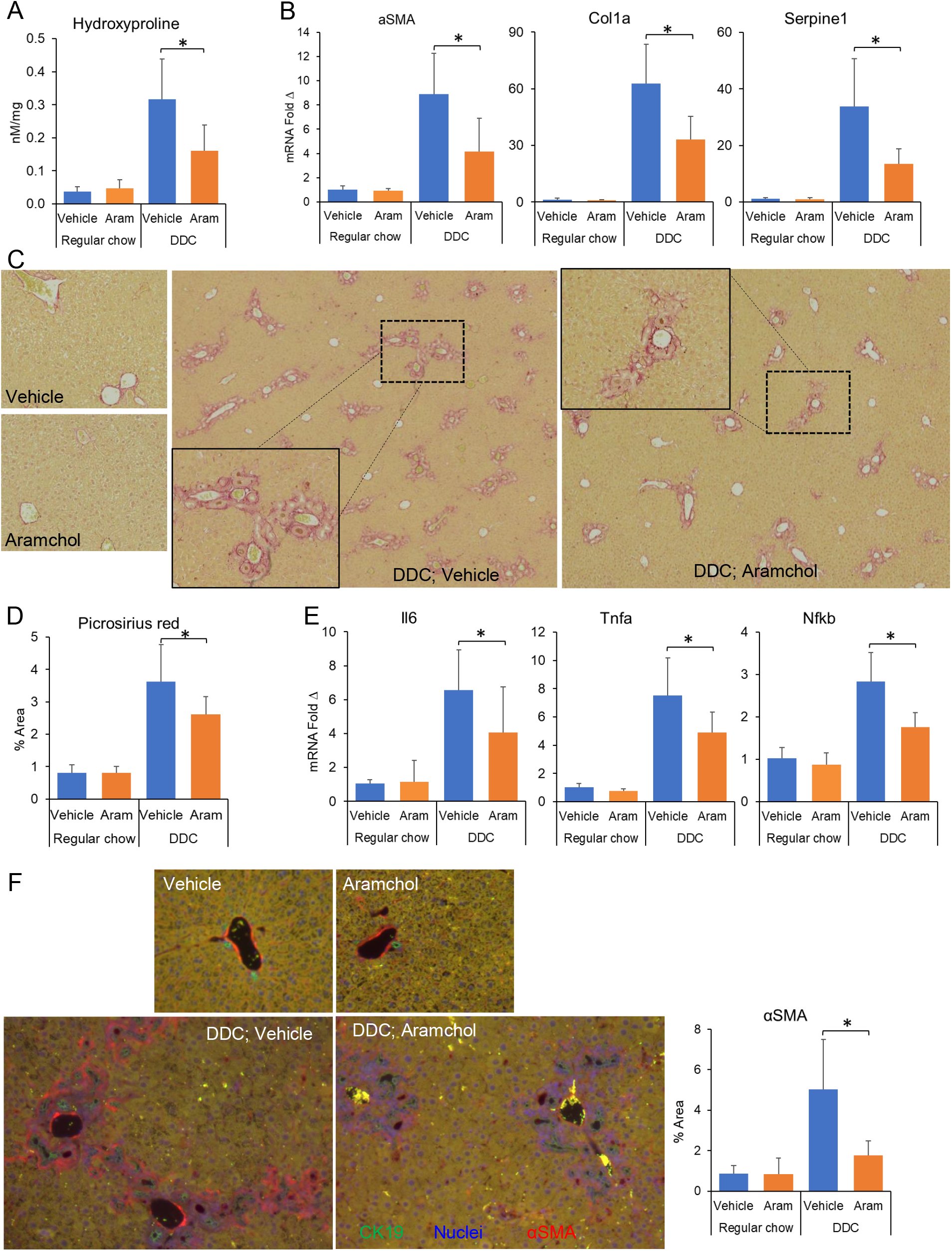
A) The increased liver content of hydroxyproline caused by the DDC diet was significantly reduced with Aramchol meglumine treatment (shortened to Aramchol in the figures) compared to vehicle only. B) qPCR analysis demonstrated Aramchol meglumine significantly inhibited the DDC diet-induced increase in ECM components, *Col1a* and *αSMA*, and fibrotic marker, *Serpine1*. C) The DDC diet-induced increased picrosirius staining of liver sections was significantly attenuated with Aramchol meglumine treatment with area quantification shown in D. E) The DDC diet-induced increased expression of inflammatory markers, *Il6, Tnfa* and *Nfkab*, were significantly inhibited with Aramchol meglumine treatment. F) The DDC diet-induced increased IF of αSMA (red) surrounding CK19 immunostained bile ducts (green) was significantly reduced with Aramchol meglumine treatment. (N = 9 mice per group, *= p<0.05).

### Aramchol inhibits the TGFβ-stimulated fibroinflammatory gene expression in cholangiocyte

We have previously demonstrated that cholangiocytes activated with TGFβ express a number of fibrosis and inflammation stimulating genes [27]. TGFβ may regulate transcriptomics by modulating cellular metabolomics including upregulation of enzymes involved in synthesizing inflammatory lipids [11, 12]. In fact, TGFβ upregulated the expression of SCD1 in retinal epithelial cells) [11-13]. Aramchol was previously shown to both downregulate SCD1 and inhibit its function [18, 19]. Using RNA-seq, we show that Aramchol broadly inhibited the TGFβ transcriptomic modulations (Fig 3A). Comparing TGFβ and Aramchol co-treated H69 cells with TGFβ only, 558 genes were significantly downregulated while 336 genes were significantly upregulated with Aramchol co-treatment (adjusted p-value <0.05). These genes were subjected to Ingenuity Pathway Analysis (IPA) to identify the most effected signaling pathways (those with the lowest p-values). Fibrosis and hepatic stellate cell activation pathways were the most downregulated, while anti-fibroinflammatory PPAR signaling was among the most upregulated pathways (Fig. 3B). These observations indicate that Aramchol attenuates the TGFβ-induced gene expression with prominent effects in downregulating fibrotic pathways while upregulating anti-fibroinflammatory signaling. Representative genes of the downregulated pathways, interleukin 6 (IL6), vascular endothelial growth factor A (VEGFA), platelet-derived growth factor B (PDGFB), and plasminogen activator inhibitor-1 (PAI-1, *SERPINE1* gene) were analyzed in transformed (H69 and MLE) and human primary PSC cholangiocytes (PSC-Cs). The TGFβ-induced increased expression was significantly inhibited by Aramchol in a dose dependent manner both at the transcript and protein level (Fig. 3C and D). In PSC-Cs, the basal high expression of these genes was markedly downregulated by Aramchol except *SERPINE1* (Fig. 4A). IL6, which was abundantly released into the cell cultured media (CCM) by PSC-C, was significantly reduced by Aramchol determined by a specific ELISA assay (Fig. 4B). Similarly, in MLEs, the TGFβ-induced expressions of *Il6, Vegfa, Pdgfb* and *Serpine1* were significantly reduced by Aramchol in a dose-dependent manner (Fig. 3C), as was the IL6 released into the CCM determined by ELISA (Fig. 4D). SCD expression was significantly reduced in PSC-Cs but not in the transformed cells (Fig. 3C and Fig. 4A). These observations indicate that Aramchol markedly inhibits the fibroinflammatory signals emanating from activated cholangiocytes consistent with previous studies in HSCs [19].

**Fig 3.**
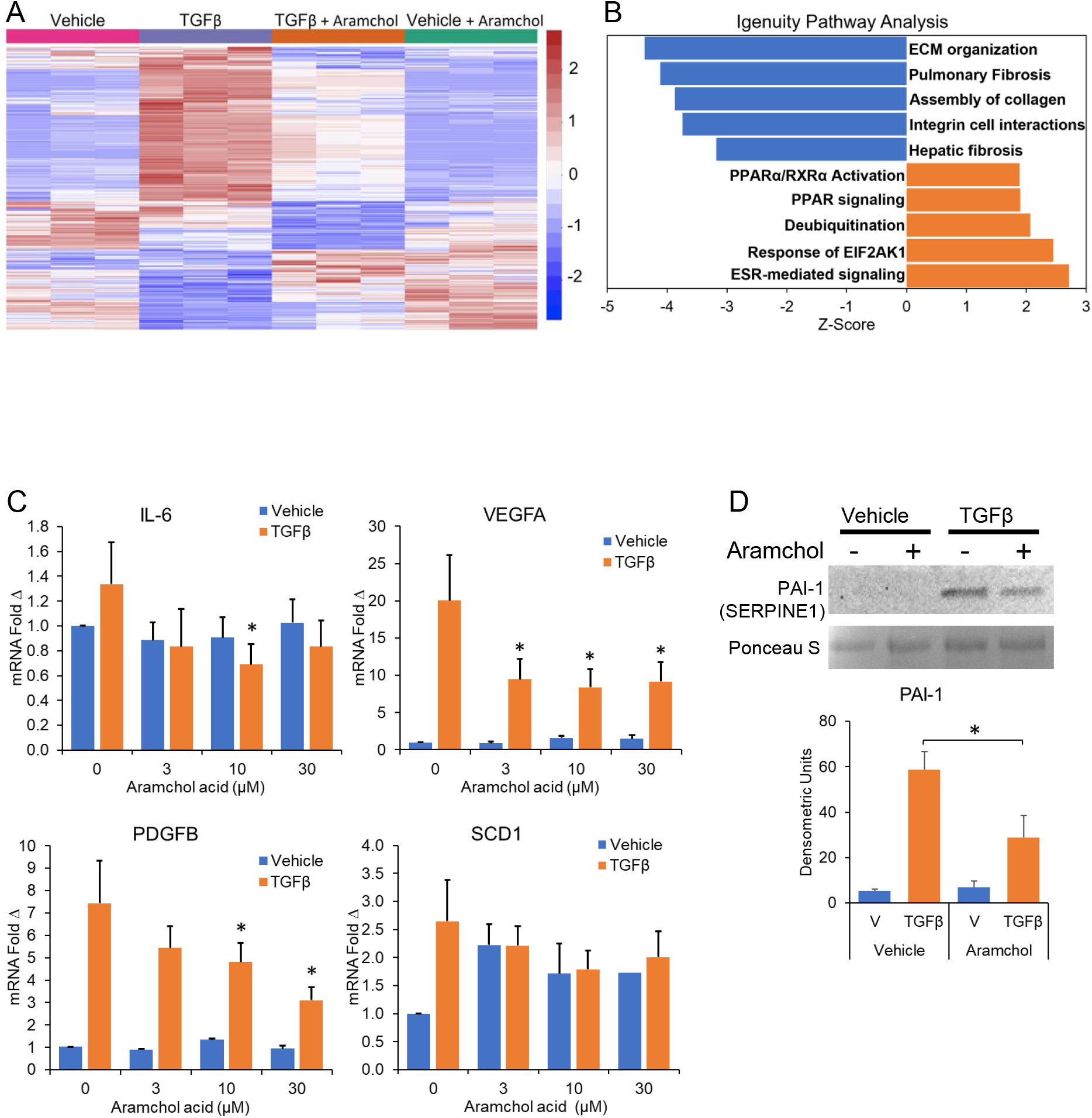
A) RNA-seq analysis of H69 cholangiocytes revealed significant modulation of multiple genes with TGFβ stimulation (columns 1 vs 2), which were significantly attenuated with Aramchol co-treatment (columns 2 vs 3). B) IPA analysis of genes that showed significant changes with Aramchol co-treatment compared to TGFβ only (columns 3 vs 2 of A) identified inhibition of hepatic fibrosis pathways (negative Z-scores) while stimulating PPAR signaling (positive Z-score) by Aramchol co-treatment. C) qPCR analysis showed that the TGFβ-induced elevated levels of *IL6, VEGFA* and *PDGFB* were significantly and dose-dependently inhibited with Aramchol co-treatment in H69 cholangiocytes. D) Western blot analysis demonstrated that the TGFβ-induced elevated level of PAI1 was significantly inhibited with Aramchol co-treatment. (N=3 at least, *= p<0.05 when compared to vehicle or TGFβ only).

**Fig 4.**
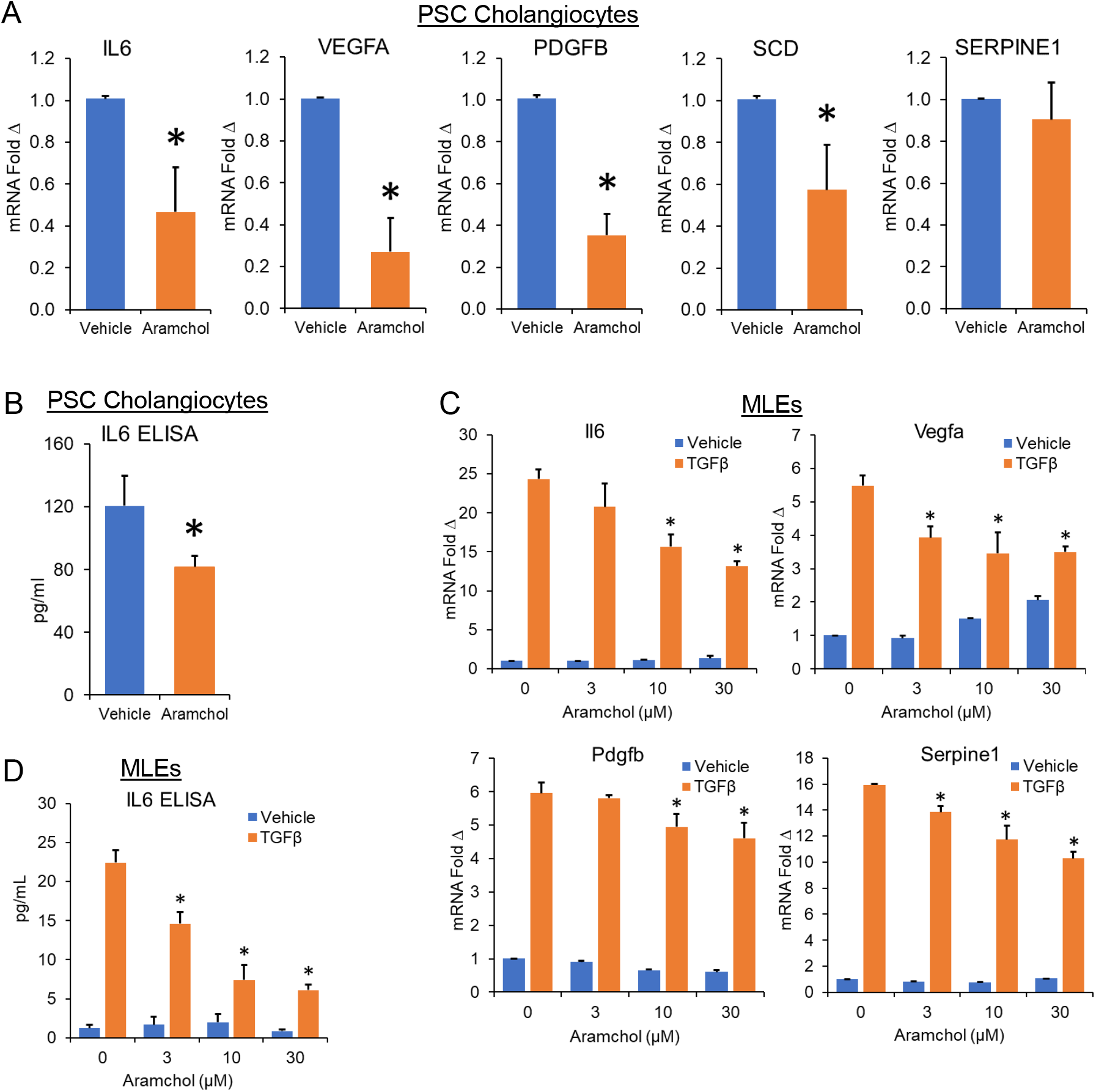
A) qPCR analysis showed that Aramchol treatment of PSC-Cs resulted in significant reductions in the expression of *IL6, VEGFA, PDGFB*, and *SCD* but not *SERPINE1*. B) ELISA analysis of PSC-C cultured media demonstrated significant reductions in IL6 level with Aramchol treatment. C) qPCR analysis showed that the TGFβ-induced elevated expression of *Il6, Vegfa, Pdgfb* and *Serpine1* were significantly and dose-dependently inhibited with Aramchol co-treatment. D) ELISA analysis showed that the TGFβ-induced elevated level of Il6 was significantly and dose-dependently inhibited with Aramchol co-treatment. (N=3 at least, *= p<0.05 when compared to vehicle or TGFβ only).

### SCD siRNA knockdown in cholangiocytes shows similar results to Aramchol treatment

Aramchol has been previously shown to downregulate markers of fibrosis in HSCs and hepatocytes by inhibiting SCD [18, 19]. Using H69 cholangiocytes, we show that SCD siRNA knockdown has similar effects to Aramchol treatment. SCD was significantly knocked down by ∼95%, which resulted in significantly attenuating the TGFβ-induced mRNA expression of *IL6, SERPINE1* and *VEGFA*, and protein level of IL6 and PAI-1 (Fig. 5). These observations indicate that the effects of Aramchol on cholangiocytes, similar to HSCs and hepatocytes, are through inhibition of SCD.

**Fig 5.**
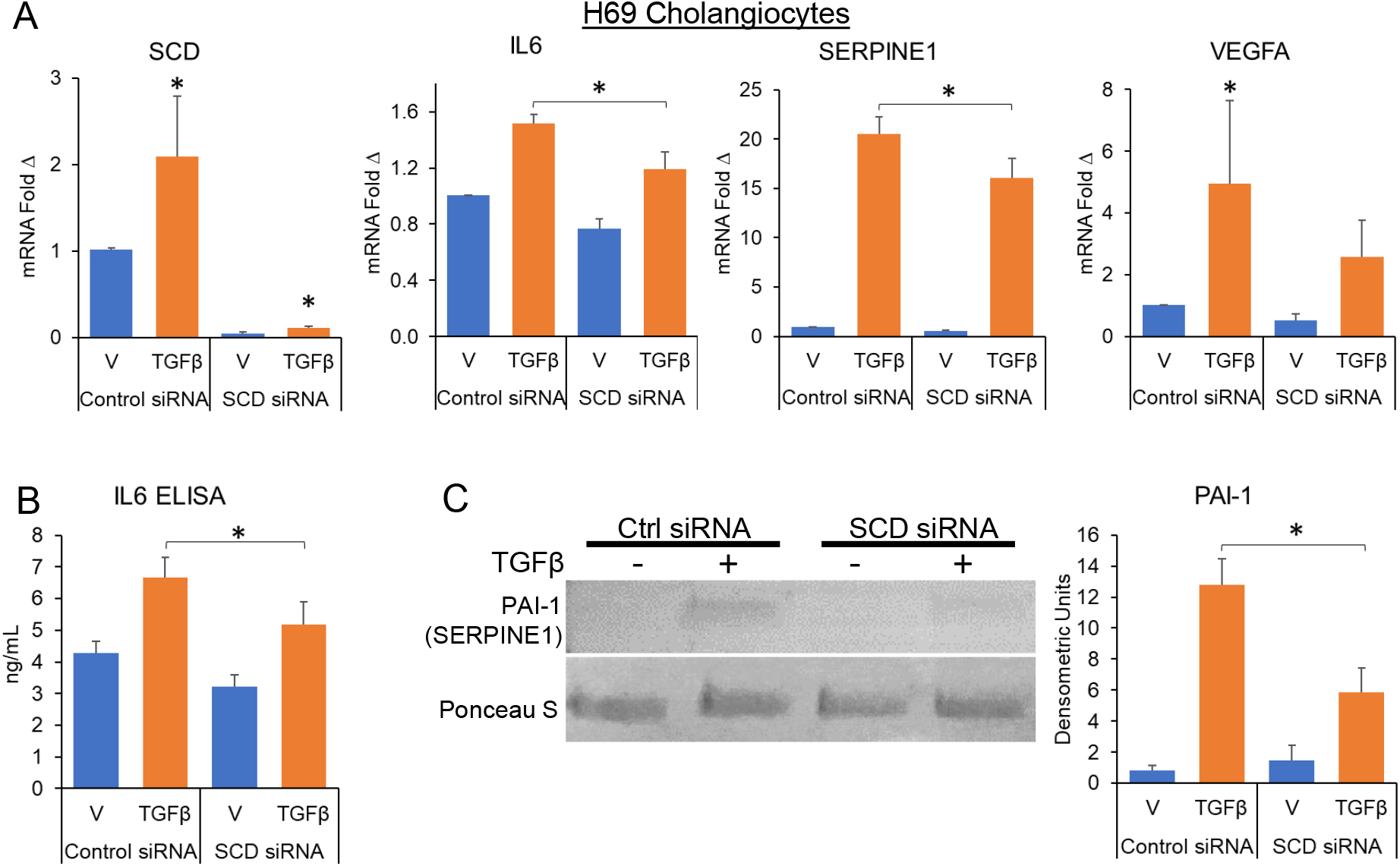
A) qPCR analysis showed increased expression of SCD with TGFβ treatment of H69 cholangiocytes, which was significantly knocked down (∼95%) with specific siRNAs. Furthermore, the TGFβ-induced elevated expression of *IL6* and *SERPINE1* were significantly inhibited with SCD siRNA knockdown. B&C) ELISA and Western blot analyses showed that the TGFβ-induced elevated expression of IL6 and PAI1 respectively in the cell cultured media were significantly inhibited with SCD siRNA knockdown. (N=3 at least, p<0.05).

### Aramchol upregulates PPARα/γ activity

TGFβ signaling inhibits PPARα/γ, while activation of PPARα/γ suppresses the fibroinflammatory signals [28, 29]. Aramchol has been previously shown to upregulate PPARγ expression in HSCs [18-20]. Here, we show that Aramchol increased the PPARα and γ binding to PPRE sites, a direct measure of PPAR activity, in TGFβ-stimulated H69 cholangiocytes (Fig. 6A). Consistently, PPARα/γ responsive genes were increased (Fig. 6B). However, PPARG expression was not increased with Aramchol (Fig. 6B). In MLEs, Aramchol significantly increased the PPARα and γ binding to PPRE sites under basal conditions, but only significantly increased the activity of PPARγ when cholangiocytes were co-stimulated with TGFβ (Fig. 6C). *Pparg* and *Ppara* expression was significantly downregulated with TGFβ, which was reversed with Aramchol (Fig. 6D). Correspondingly, the TGFβ-induced suppression of PPAR-responsive genes was also significantly reversed (Fig. 6D).

**Fig 6.**
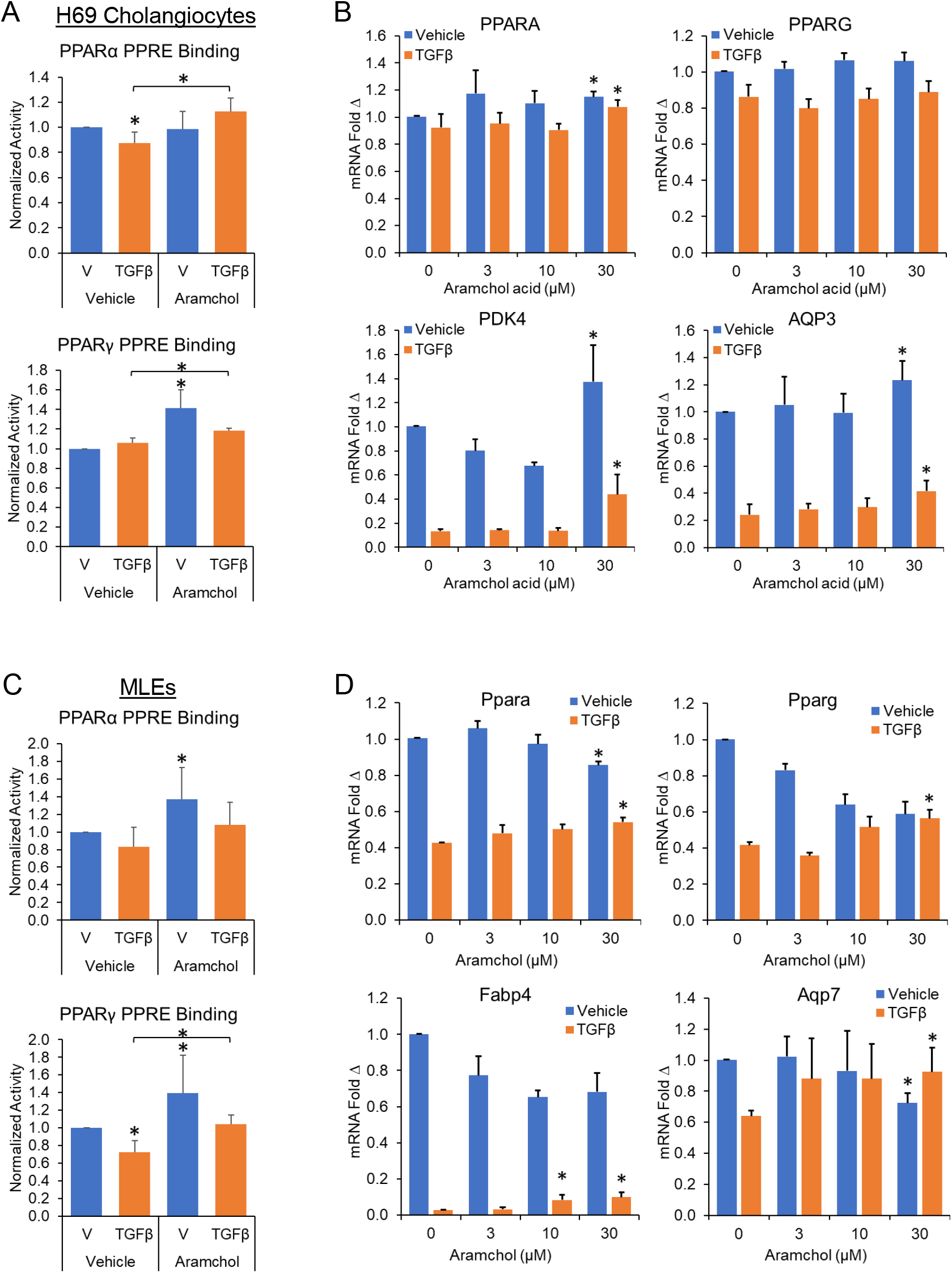
A) Using an ELISA system with immobilized DNA containing PPRE sites and PPAR specific antibodies, PPARα binding to PPRE sites was significantly reduced with TGFβ treatment, which was significantly reversed with Aramchol co-treatment in H69 lysates. PPARγ binding of PPRE sites was also increased with Aramchol treatment. B). *PPARA* mRNA expression was significantly increased with Aramchol but not *PPARG*. PPARα/γ responsive genes, *PDK4* and *AQP3*, were significantly reduced with TGFβ, which was partially reversed with Aramchol co-treatment. C) In MLEs, Aramchol significantly increased the PPARα and γ binding to PPRE sites under basal conditions, but only significantly increased the PPRE binding of PPARγ with co-treatment of TGFβ. D) *Pparg* and *Ppara* expression was significantly downregulated with TGFβ, which was partially but significantly reversed with Aramchol. The TGFβ-induced suppression of PPAR-responsive genes, *Fabp4* and *Aqp7*, was also significantly reversed. (N=3 at least, *= p<0.05 when compared to vehicle or TGFβ only).

**Fig 7.**
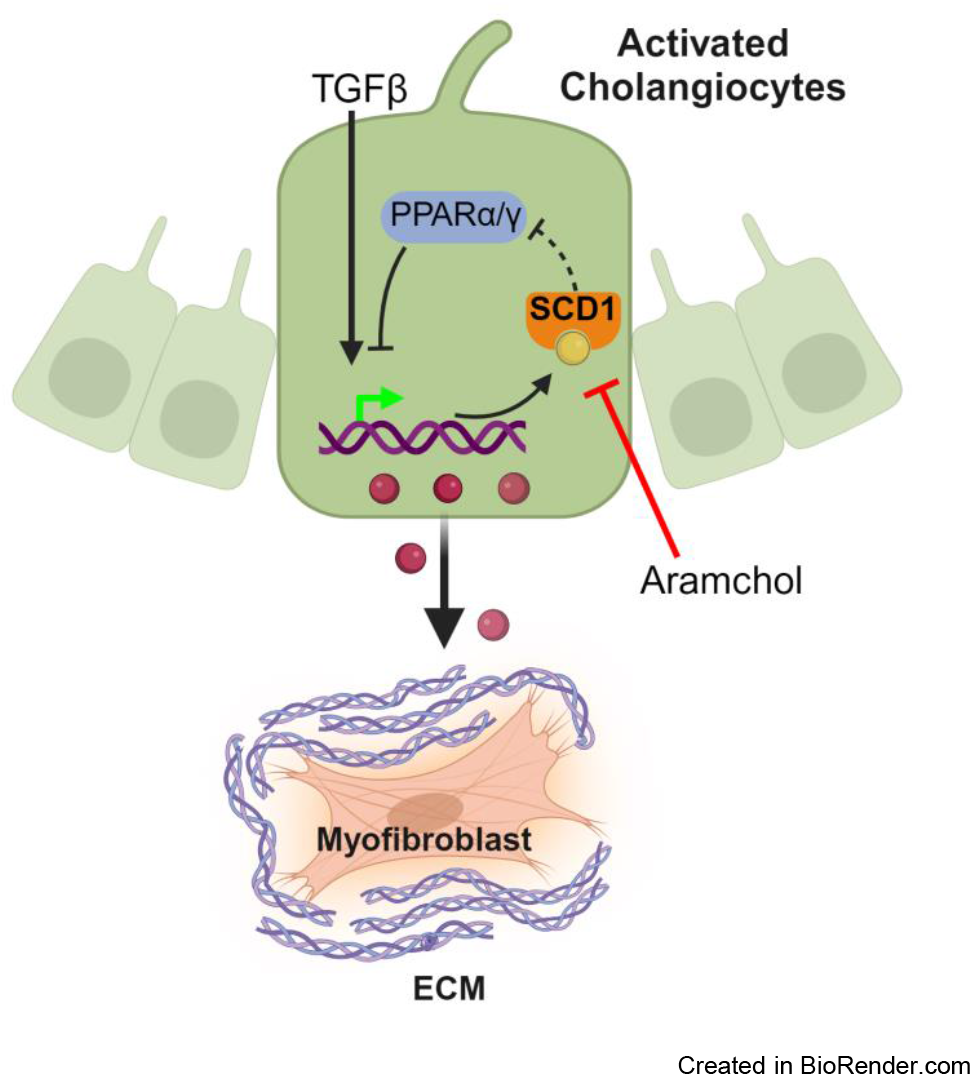
TGFβ signaling upregulates SCD1, which inhibits PPARα/γ, allowing for the expression of fibroinflammatory signals in cholangiocytes. Aramchol inhibition of SCD1 allows increased PPARα/γ expression and activity, which inhibits the TGFβ-induced expression of fibroinflammatory signals.

## DISCUSSION

Biliary fibrosis is the predominant process of fibroinflammatory cholestatic liver diseases such as PSC and PBC, where morbidity and mortality closely correlate with biliary fibrosis. There are currently no treatments for biliary fibrosis, which severely limits therapeutic options for these diseases. Central to the pathogenesis of biliary fibrosis are cholangiocytes that serve as a ‘hub’ of signaling [30]. Injury and inflammatory signals activate cholangiocytes to become highly secretory. Activated cholangiocytes interact with immune cells and myofibroblasts to initiate and propagate biliary fibrosis in the setting of chronic disease [8]. However, the molecular mechanisms of activated cholangiocytes that allow for its secretory phenotype and paracrine interactions are not completely elucidated.

In this study, we showed that Aramchol meglumine significantly attenuated biliary fibrosis in two mouse models of cholestatic injury and fibrosis, the Mdr2^-/-^ and the DDC diet model. This effect was demonstrated by reductions in both measures of ECM synthesis (mRNA expression of ECM components in the liver) and the collagen composition of the liver (picrosirius red staining and hydroxyproline content). Interestingly, these reductions in fibrosis were not accompanied by a significant change in liver injury markers such as ALT, AST and ALP (Supplementary Fig 1 and 2). These observations suggest that Aramchol meglumine does not have a significant effect on bile toxicity or cholestasis-induced liver injury. This is unlike bile acid therapies such as ursodeoxycholic acid with direct reduction of bile and cholestasis toxicity compared to this fatty acid-bile acid conjugate [31, 32]. Rather, the effects of Aramchol are likely mediated through cellular signaling post-injury, which can be combined with toxicity-reducing modalities. We hypothesized that the attenuation of biliary fibrosis by Aramchol results from its effects on the cholangiocyte signaling. Accordingly, we tested Aramchol’s effects on the inhibition of pro-fibroinflammatory signals emanating from cholangiocytes. Cholangiocytes activated by TGFβ release a number of inflammatory and fibrogenic stimuli. These signals mediate the interactions and activation of immune cells and myofibroblast, which collectively propagate inflammation and fibrosis. Inhibiting the signaling pathways that stimulate these signals attenuates biliary fibrosis [30]. Consistently, Aramchol co-treatment of cholangiocytes led to significant attenuation of TGFβ-induced fibroinflammatory signals. This is hypothesized to occur through at least two interdependent modalities: inhibition of SCD1 and activation of PPARα and PPARγ.

TGFβ upregulates SCD1 in epithelial cells, which may be partially responsible for TGFβ-induced downstream effects [11-13]. Aramchol has been previously shown to inhibit SCD1 in hepatocytes and HSCs [18-20]. As further evidence, we show that SCD1 siRNA knockdown in cholangiocytes produced similar effects to Aramchol treatment on TGFβ-induced gene expression. These observations support the hypothesis that Aramchol attenuates the fibroinflammatory effects of TGFβ by inhibiting SCD1 in cholangiocytes.

PPAR signaling promotes anti-fibroinflammatory pathways in part by inhibiting TGFβ signaling. On the other end, TGFβ inhibits PPARγ to allow for its fibrogenic signaling [28, 29]. Previous studies have shown that SCD1 activity can inhibit PPARγ [11-13]. In HSCs, Aramchol treatment or SCD1 siRNA knockdown resulted in upregulation of PPARγ [19]. Therefore, the TGFβ inhibition of PPARγ may occur through upregulation of SCD1 expression and activity. Consistently, Aramchol treatment of cholangiocytes upregulates PPARα/γ expression and activity. PPAR activity was determined by both measurement of PPRE binding and assessment of PPAR-responsive genes, which were both upregulated with Aramchol co-treatment. These observations support the interplay of TGFβ, SCD1 and PPARγ in cholangiocyte-mediated promotion of biliary fibroinflammation.

The effects of Aramchol may extend beyond the TGFβ signaling pathway given the complex interactions of this pathway with several others and the involvement of SCD1 in other pathways [33, 34]. In fact, SCD1 is implicated in Wnt signaling to promote HSC activation and hepatocyte tumorigenesis [35]. The canonical Wnt pathway is well-studied in biliary fibrosis and known to overlap with TGFβ signaling [33].

The attenuation of fibrosis in our *in vivo* models may also be in part due to Aramchol directly inhibiting hepatic myofibroblast ECM synthesis. Indeed, *in vitro* studies of Aramchol have demonstrated inhibition of HSC production of ECM components [19]. The beneficial effects of Aramchol may extend to other cell compartments involved in biliary fibrosis, including the cells of the immune system [30, 36, 37]. Indeed, SCD1 impairs the reparative phenotype of macrophages while stimulating an inflammatory phenotype in models of multiple sclerosis [38]. Macrophages have important roles in the pathogenesis of biliary fibrosis, both reparative (resident Kupffer cells) and deleterious (monocyte derived) [39, 40]. SCD1 knockdown resulted in increased regulatory T-cell differentiation by activation of PPARγ and reduction of autoimmunity [41]. However, any direct effects of Aramchol or SCD1 on immune cells and their activation or differentiation has not been shown in detail in models of biliary fibrosis. To distinguish the effects of Aramchol on various cell mediators involved in the pathogenesis of biliary fibrosis would require transgenic mouse models to specifically knockdown Aramchol target, SCD1, in these cell compartments.

Aramchol is well-studied in MASLD with promising results. It reduces steatohepatitis and fibrosis in mouse models of MASLD [20]. In a randomized, double-blind, placebo-controlled phase IIb trial and an open label extension of a phase III study, Aramchol demonstrated significantly increased resolution of steatohepatitis and improvement in fibrosis while decreasing liver injury markers such as serum ALT [21, 22]. Importantly, Aramchol was safe, well tolerated and without an imbalance in adverse events compared to the placebo arm. A similar safety level was observed in a meta-analysis of 3 clinical trials [42].

Taken together, Aramchol attenuates biliary fibrosis in two mouse models of biliary fibrosis along with anti-fibrotic effects in cholangiocytes, myofibroblasts and hepatocytes. These observations combined with its excellent clinical trial safety data provide the rationale for further clinical studies of Aramchol in patients with biliary fibrosis, in particular PSC, where treatments are desperately needed.

## Acknowledgements

This study was funded by the manufacturer of Aramchol, Galmed Pharmaceuticals. Additional funding came from the Stravitz-Sanyal Institute for Liver Disease and Metabolic Health and the Department of Internal Medicine Pilot Fund, VCU. The data included in this study was generated at the Genomics Core facility at VCU. Services in support of the research project were provided by the VCU Massey Comprehensive Cancer Center Bioinformatics Shared Resource. Massey is supported, in part, with funding from NIH-NCI Cancer Center Support Grant P30 CA016059. Services and products in support of the research project were generated by the Lipidomics and Metabolomics Shared Resource, supported, in part, with funding from NIH-NCI Cancer Center Support Grant P30 CA016059. We sincerely thank Dr. Daniel Goldenberg (Hadassah-Hebrew University Medical Center, Jerusalem, Isreal) for providing the Mdr2^-/-^ mice and Dr. Nicholas LaRusso (Mayo Clinic, Rochester, MN, USA) for providing PSC-Cs.

## Figure legends

**Supplementary Table 1.**
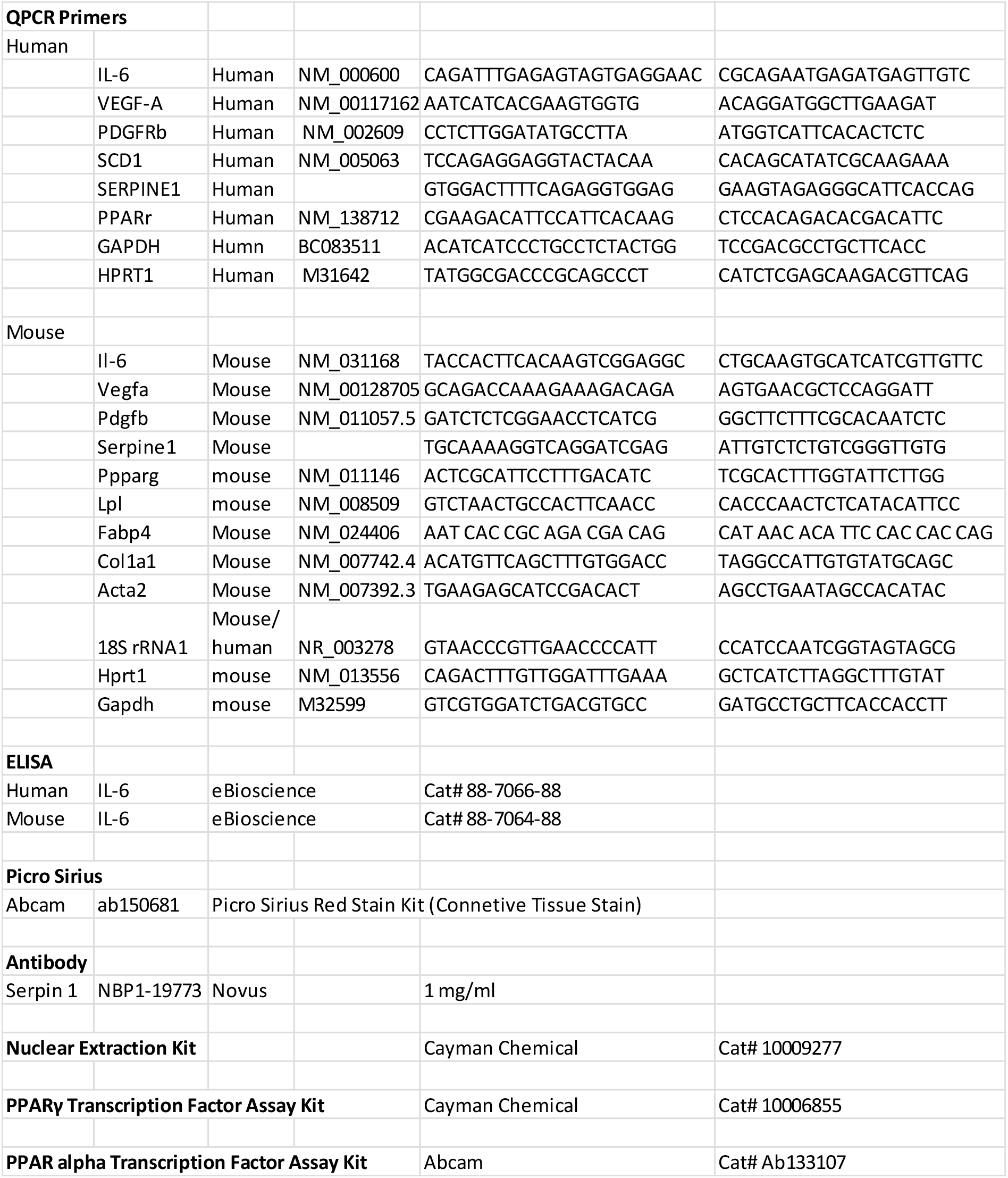

**Supplementary Fig 1.**
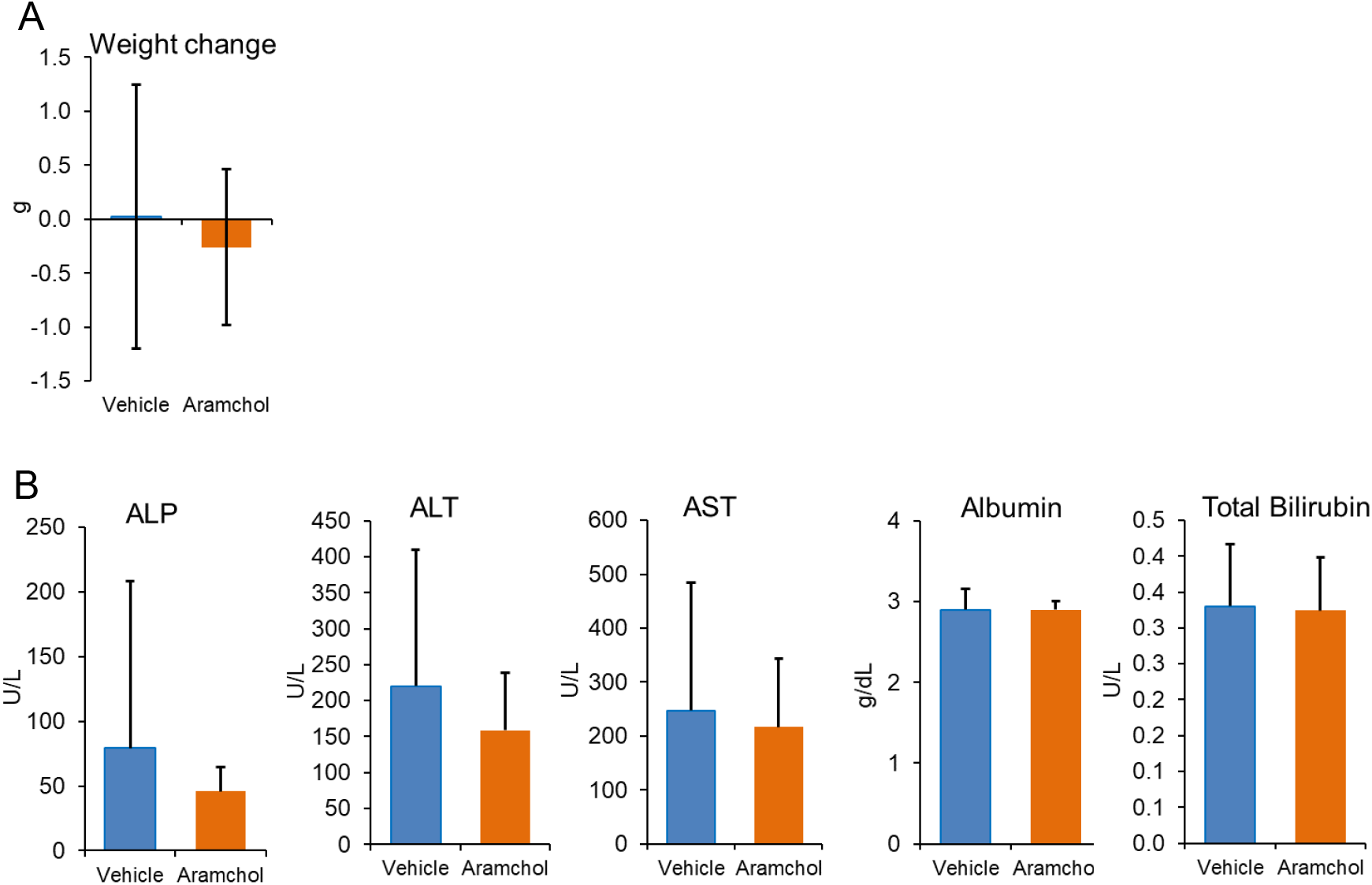
A) Weight change at the end of the experiment compared to baseline was not significantly different between Aramchol and vehicle only treated mice. B) Serum hepatic injury markers (alkaline phosphatase (ALP), alanine aminotransferase (ALT), and aspartate aminotransferase (AST)) or hepatic function tests (total bilirubin and albumin) were not significantly different between Aramchol and vehicle only treated mice. (N=9 mice per group).

Supplementary Fig 1. A) The DDC diet resulted in significant weight loss at the end of the 3-week experiment, but there were no significant differences between Aramchol and vehicle treated mice. B) The DDC diet resulted in a significant rise in serum hepatic injury markers (alkaline phosphatase (ALP), alanine aminotransferase (ALT), and aspartate aminotransferase (AST) and measure of biliary obstruction (total bilirubin) but not albumin. There were no significant differences in these labs between Aramchol and vehicle only treated mice. (N=9 mice per group).

**Supplementary Fig 2.**
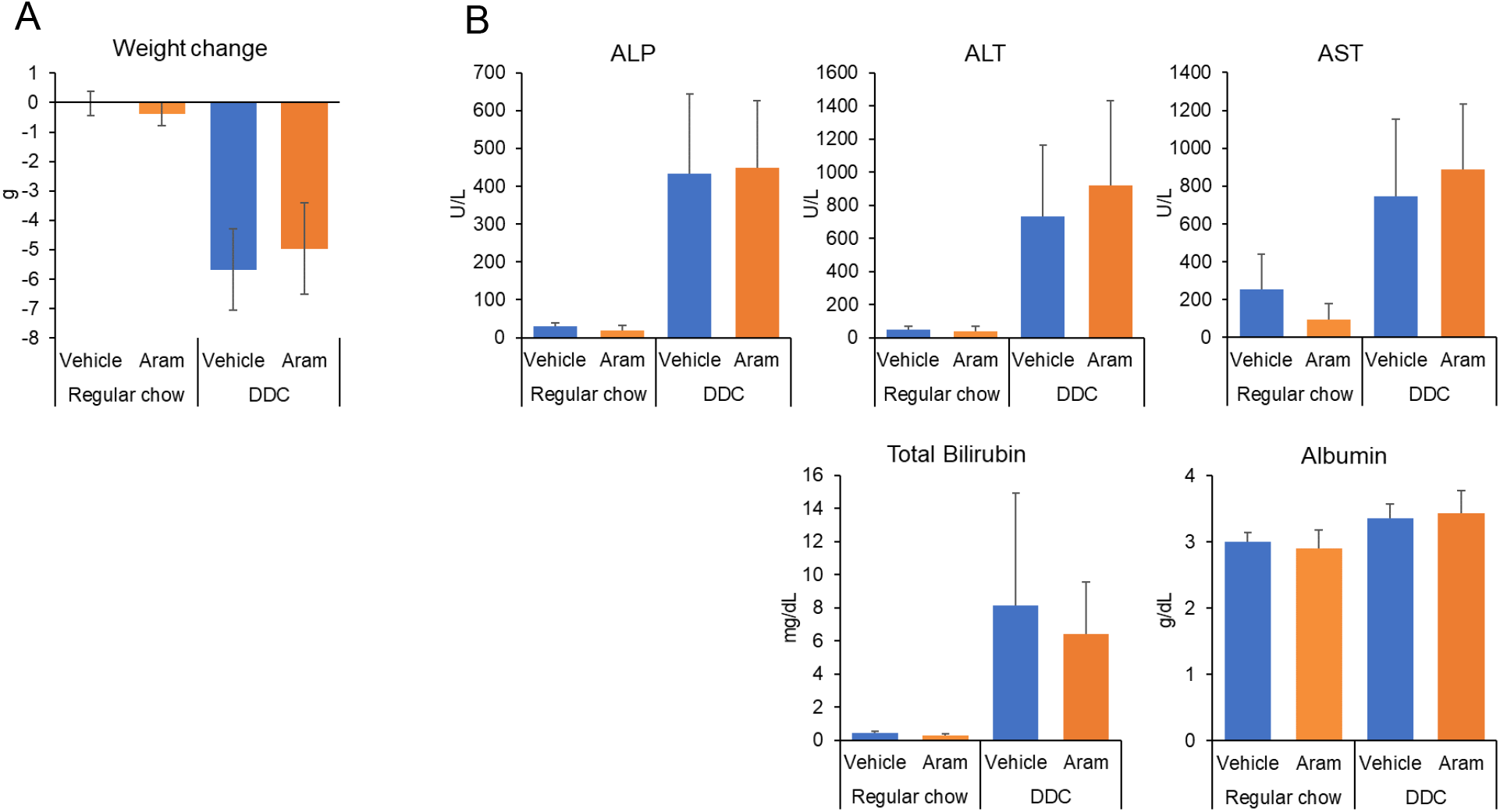

## Notes

### Summary of Updates

The text of the manuscript was edited to clarify the results, methodologies and discussion.

